# Female mice are resilient to age-related decline of substantia nigra dopamine neuron firing parameters

**DOI:** 10.1101/621680

**Authors:** Rebecca D. Howell, Sergio Dominguez-Lopez, Sarah Ocañas, Willard M. Freeman, Michael J. Beckstead

## Abstract

The degeneration of substantia nigra (SN) dopamine neurons is a central feature in the pathology associated with Parkinson’s disease, which is characterized by progressive loss of motor and cognitive functions. The largest risk factors for Parkinson’s disease are age and sex; most cases occur after age 60 and males have nearly twice the incidence as females. While much research in Parkinson’s has focused on the mechanisms underlying dopamine neuron degeneration, very little work has considered the influence of these two risk factors to disease risk and presentation. In this work, we performed whole cell patch clamp recordings in brain slices to study the alterations in intrinsic firing properties of single dopamine neurons in C57BL/6 mice across ages and between sexes. We observed a progressive decline in dopamine neuron firing activity in males by 18 months of age, while dopamine neurons from females remained largely unaffected. A semiquantitative analysis of midbrain dopamine neuron populations revealed a slight decrease only in substantia nigra dopamine neurons in males, while females did not change. This was also accompanied by increases in the expression of genes that have been linked to Parkinson’s including PTEN-induced kinase 1 (PINK1) in both males and females, and the ubiquitin ligase parkin, primarily in the substantia nigra of males. These impairments in dopamine neuron function in males may represent a vulnerability to further insults that could predispose these cells to neurodegenerative diseases such as in Parkinson’s.

## INTRODUCTION

Midbrain dopamine neurotransmission is involved in a wide array of functions including motor control, motivation, reward, and cognitive processes (Crocker, 1997; Wise & Rompre, 1989). The dopaminergic cell bodies of the substantia nigra (SN) and the ventral tegmental area (VTA) form the starting points for the nigrostriatal and mesolimbic pathways, respectively (Fuxe, Hokfelt, Olson, & Ungerstedt, 1977). Degeneration of dopamine neurons in the SN is a hallmark feature of the pathology of Parkinson’s disease and is associated with a progressive loss of the ability to initiate movement (Hornykiewicz, 1975). Parkinson’s disease is the result of multiple complex gene/environment interactions (Duda, Potschke, & Liss, 2016) and recent studies suggest that the incidence may be increasing (Savica, Grossardt, Bower, Ahlskog, & Rocca, 2016). Two of the most commonly observed risk factors for the development of Parkinson’s are age and sex (Gillies, Pienaar, Vohra, & Qamhawi, 2014). Most reports show an increase in the incidence of Parkinson’s disease after the age of 60 (de Lau & Breteler, 2006) and many also indicate a 1.5 to 2 times greater incidence in males compared with females (Baldereschi et al., 2000; Moisan et al., 2016). Despite the awareness of these two risk factors in the clinical literature, little basic research has considered the effects of age and sex on disease development and presentation.

Previous work in our lab identified a decrease in spontaneous firing frequency and a lack of pacemaker firing fidelity in dopamine neurons recorded in slices from male mice over 25 months of age (Branch, Sharma, & Beckstead, 2014). Dopamine neurons exhibit rhythmic, spontaneous pacemaking activity in brain slices (Grace & Onn, 1989), which is the result of the coordinated activity of at least a dozen ion channels, calcium homeostasis, and energy metabolism (Duda et al., 2016). The maintenance of continuous activity renders dopamine neurons highly vulnerable to oxidative damage, which is thought to be a possible mechanism for their degeneration in Parkinson’s disease (Duda et al., 2016). Dysfunction in firing suggests deficits in the health of the cells and possibly impairment of the nigrostriatal circuit with consequences for motor function (Branch et al., 2016).

In the present study we sought to perform a full characterization of dopamine firing deficits, determine when during the mouse lifespan dopamine cell deficits appear, and explore any potential effects of sex. To do this, we performed current clamp recordings of SN dopamine neurons across four age groups in both male and female C57Bl/6 mice. The age groups tested were 4, 12, 18, and 24-30 months, which correspond roughly to human ages 25, 42, 56, and 69-81 (Fox, 2007). We found that dopamine neurons from females show few obvious decrements with age, while neurons from males exhibited firing deficits by 18 months. In order to better understand the implications of these results, we then complemented these findings with a semi-quantitative immunocytochemistry analysis of midbrain dopamine neurons in young and old, male and female animals as well as gene expression and spontaneous locomotion studies.

## RESULTS

### Dopamine neuron pacemaking becomes disrupted in males, but not females, by 18 months of age

Dopamine neurons were identified in the ventral half of the SN of coronal midbrain slices based on their large cell bodies, characteristic slow firing rate (<7Hz) (Grace & Onn, 1989) and the presence of a large voltage sag as a result of the hyperpolarization-activated cation current, I_H_ (Dufour, Woodhouse, Amendola, & Goaillard, 2014). Recorded cells were also filled with neurobiotin to confirm their dopaminergic identity through *post hoc* immunostaining for tyrosine hydroxylase (TH), the rate limiting enzyme of dopamine synthesis (Dufour et al., 2014; Fig 1A). Dopamine neurons were recorded from male and female mice across four age groups: 4 months (n=21 cells from 6 male mice; n=23 from 3 females), 12 months (n=20 from 8 males; n=17 from 6 females), 18 months (n=21 from 4 males; n=19 from 4 females) and 24-30 months (n=31 from 10 males; n=26 from 7 females). A range of spontaneous firing frequencies were recorded in current clamp for at least 2 minutes each, across all ages in dopamine neurons from males (from 0.6 to 4.425 Hz) and females (0.76 to 5.3 Hz). An age-sex interaction was detected in firing frequency (F_3,170_=2.707, p=0.047; Fig 1B,C), with males showing lower firing frequencies than females at ages 18 months and older. This decrease in firing rate in males was more pronounced at 18 months (33.96% decrease) than at 24-30 months (16.53% decrease).

**Figure 1.**
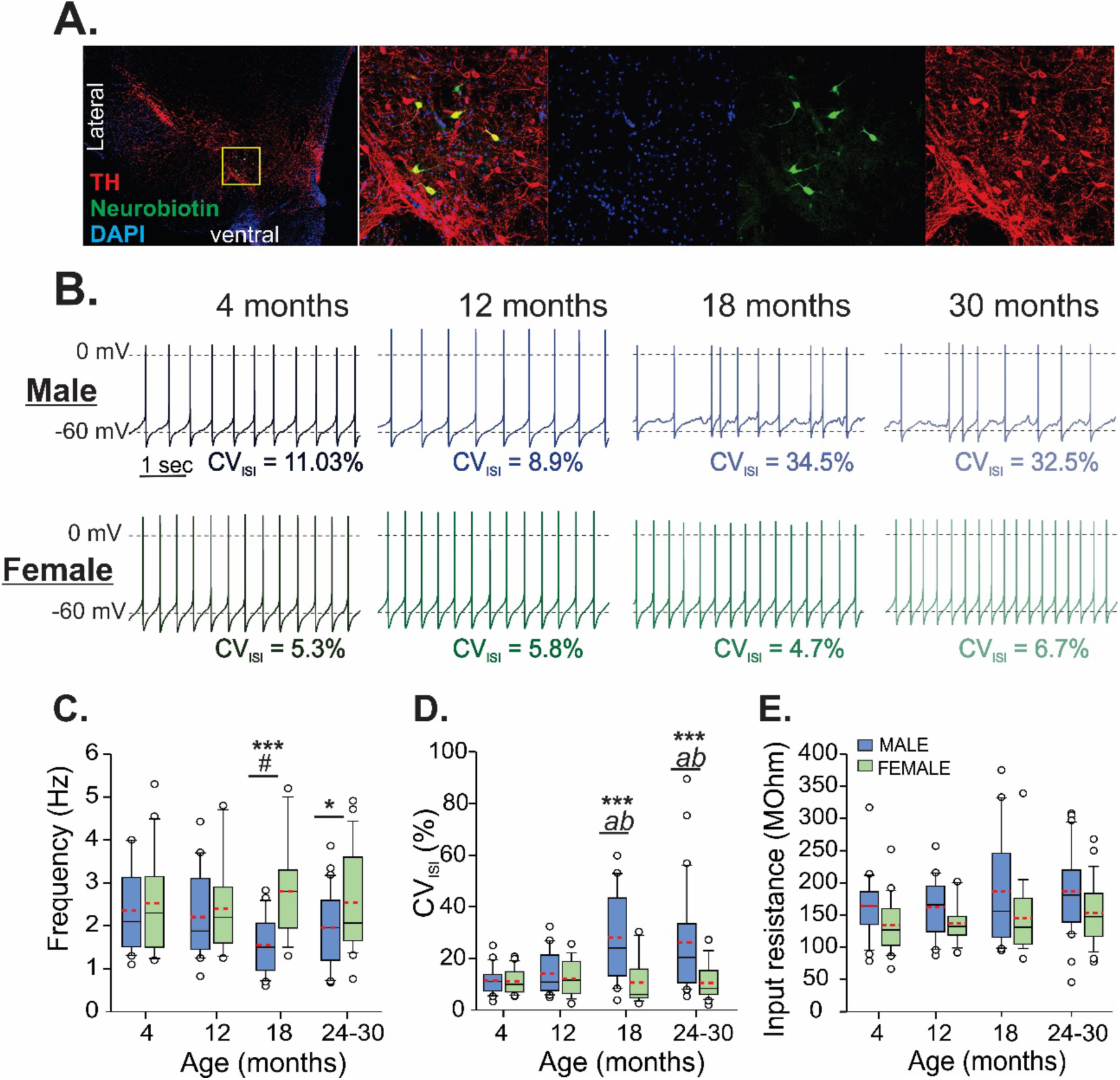
Evolution of spontaneous firing activity across ages. A. Images from a recorded slice immunolabeled for tyrosine hydroxylase (TH, a marker for DA neurons) and neurobiotin (a tracer added to the patch pipette). Far left, image of the midbrain dopamine region from the whole slice (a bisected coronal section). Images to the right show zoom in, from yellow box, of labeling for TH (red), neurobiotin (green), DAPI counterstain (blue). B. Representative recordings of spontaneous firing in current clamp from cells at 4, 12, 18, and 30 months from males (top, blue) and females (bottom, green). Box and whisker plots show firing rate (C), coefficient of variation for the interspike interval (CV_ISI_) (D), and input resistance (E). Red dashed lines indicate means. # - 18 month males differ from 4 month males, p=0.054. Males differ from females, *p<0.05, ***p<0.001; *a -* 18 or 24-30 month males differ from 4 month males, p<0.001; *b -* 18 or 24-30 month males differ from 12 month males p≤0.005

The coefficient of variation of the inter-spike interval (CV_ISI_) in spontaneous firing was measured during the most stable 30 seconds of firing of the minutes-long recordings from each cell, to assess the pacemaker function. An age-sex interaction was detected (F_3,170_=6.247, p<0.001; Fig 1E), with males showing an increase in variability of firing by 18 months that remained through 24-30 months. Although the average CV_ISI_ was much higher in males ages 18 months and older, the range of data (3.83% to 89.48%) suggest that there was a population of dopamine cells present that maintained regular pacemaking ability. Around a third of the total cells recorded from males ages 18 to 30 months had a CV_ISI_ less than 15% (23.8% of cells from 18 months and 38.7% of cells from 24 to 30 months; Fig S2). Those cells came from just half of the animals in each of the 18 and 24-30 month age groups. Of those animals, 30.95±1.94% of the cells had low CV_ISI_ in the 18-month-old group and 57.67±10.16% of the cells from the 24-30 month old group. This is consistent with substantial animal-to-animal variability in the oldest age group. Females did not show any change across age and still maintained low variability in pacemaking at ages 18 months and older (1.95% to 28.94%), with 73.33% of cells having a CV_ISI_ below 15% (Fig S2). No change in input resistance with age was observed, as previously reported (Branch et al., 2014), however, dopamine neurons from females overall showed lower input resistance compared with males (F_1,170_=14.453, p<0.001; Fig 1C).

### Action potential waveforms show divergent effects across sex and age

To further assess the underlying properties of the dopamine firing activity in males and females across age, the waveforms of the action potentials during spontaneous firing activity were averaged and analyzed for the following parameters: spike height (measured from peak to trough), half width (the duration measured at the midpoint of the spike) and afterhyperpolarization potential (AHP; raw value taken at the trough). The action potential threshold and maximum rise and decay velocities were also measured by using phase plane plots of the action potential averaged for each cell. Age-sex interactions were detected for spike height (F_3,170_=3.207, p=0.025; Fig 2C), half width (F_3,170_=3.351, p=0.02), AHP (F_3,170_=3.467, p=0.018), and maximum rise velocity (F_3,170_=5.344, p=0.002). Across the four age groups, males showed a biphasic evolution in spike height, half width, and maximum rise and decay velocities. Between 4 and 12 months across all four parameters there was an increase in spike height and rise/decay velocities and a narrowing of the half width, while after 12 months there was a steady decline in spike height, velocities and increasing half width. In contrast, females showed a more linear trajectory in these parameters with age, starting at 4 months with much higher spike heights, velocities, and narrower half widths than males, which progressively declined with age. For maximum decay velocity, only main effects of age and sex were detected (age: F_3,170_=6.778, p<0.001; sex: F_1,170_=12.346, p<0.001), whereas females showed overall higher velocities than males and both sexes showed significant decreases in velocities by 18 months. The AHP remained steady across ages in males and females until decreasing at 24-30 months in males only. No differences were detected in action potential thresholds across ages and between sexes.

**Figure 2.**
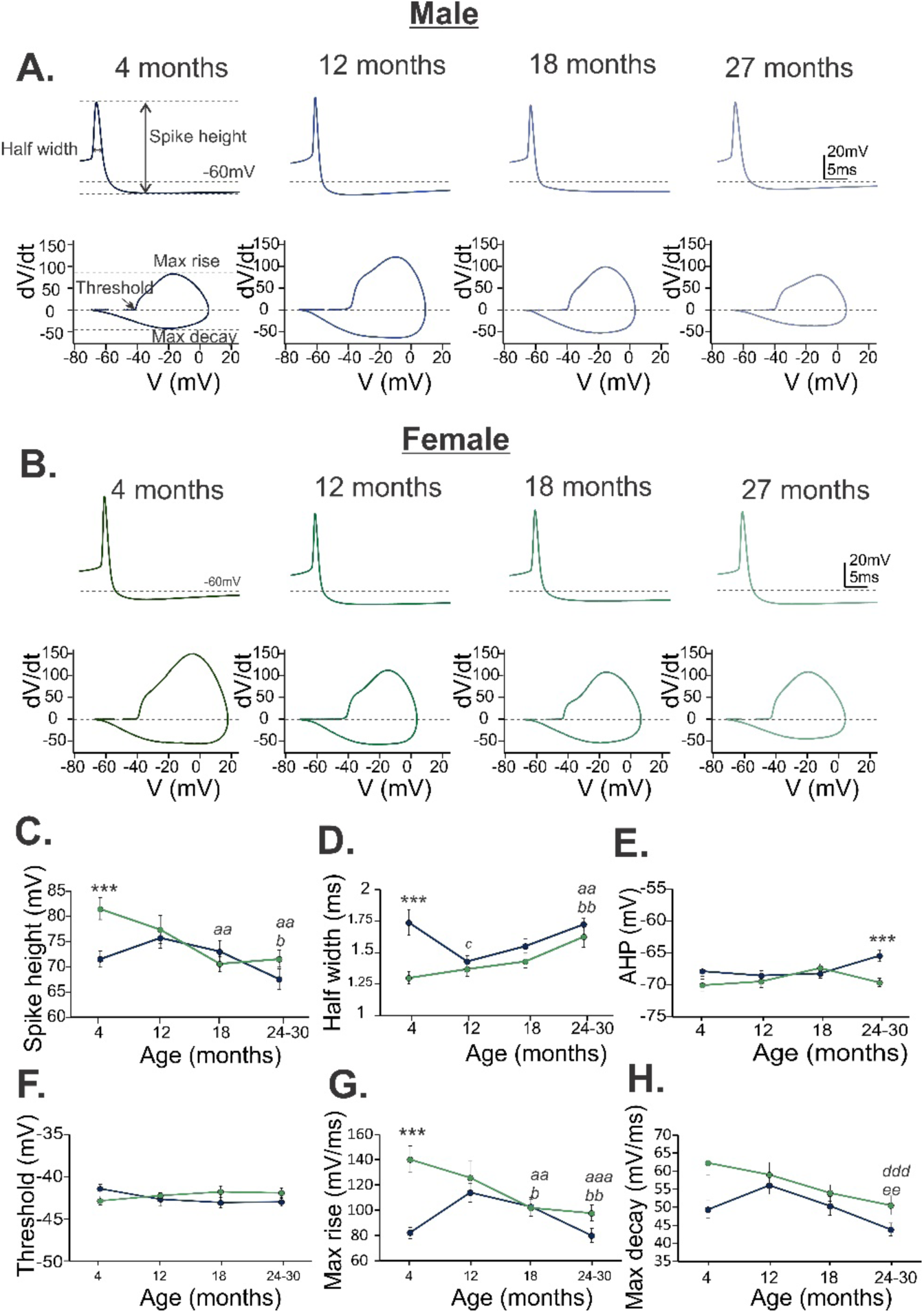
Differential effects of age on the action potential waveform in males and females. A. Top, representative averages of action potentials from cells in each of the 4 age groups of males (blue). Bottom, phase plane plots of the representative action potential traces. Parameters measured are indicated in the 4-month-old action potential and phase plane plot on the left. B. Representative action potential averages (top) and phase plane plots (bottom) from cells in each of the 4 age groups of females (green). Males (blue) and females (green) are compared in spike height (C), half width (D), AHP (E), action potential threshold (F), maximum rise velocity (G) and maximum decay velocity (H). Bonferroni corrected t-test: ***Males and females differ, p<0.001; Females 18 months or 24-30 months compared with 4 months, *aa* – p<0.005, *aaa* – p<0.001; Males 24-30 months compared with 12 months, *b* – p<0.05, bb – p<0.01; Males 12 months compared with 4 months, *c* – p<0.02; Age group 24-30 months compared with 12 months, *ddd* – p<0.001; Age group 24-30 months compared with 4 months, *ee* – p=0.003.

### Dopamine neurons from males, but not females, are more prone to depolarization block by 18 months of age

The membrane excitability of neurons was measured through frequency-current (F-I) curves in response to increasing step current injections of 20pA (20 to 400pA total). Dopamine neurons were recorded across four age groups including 4 months (n=24 from 9 males; n=21 from 3 females), 12 months (n=17 from 7 males; n=18 from 7 females); 18 months (n=22 from 7 males; n=18 from 5 females), and 24-30 months (n=29 from 9 males; n=25 from 10 females). The initial frequency was determined from the first three action potentials. In males and females, no differences were detected for initial frequency between ages nor were any differences between males and females across ages (Fig 3E).

**Figure 3.**
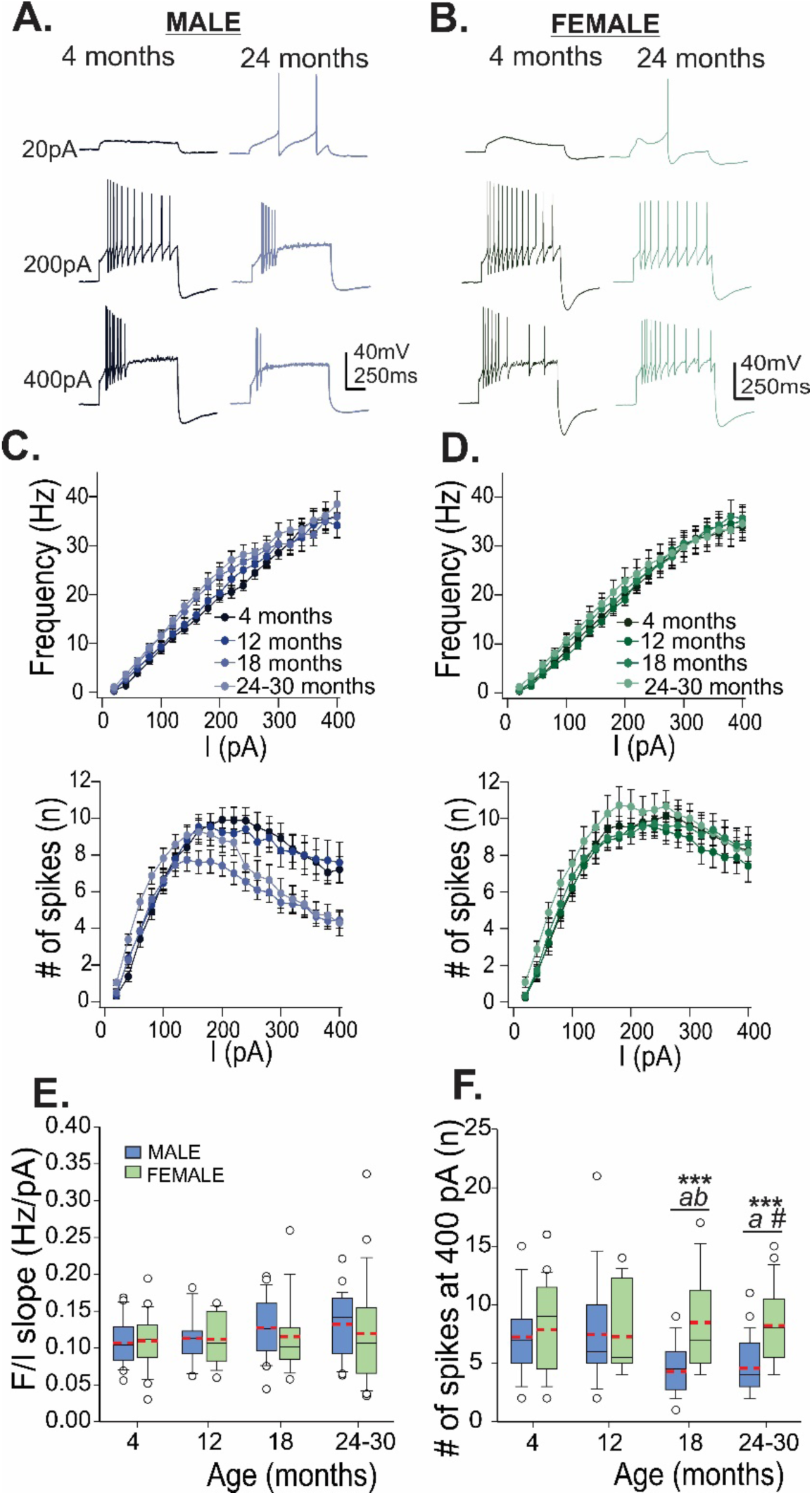
Membrane excitability across ages. Representative traces of current step injections at 20, 200, and 400 pA from males (A; blue; 4 months left, 24 months right) and females (B; green; 4 months left, 24 months right). F-I curves (top) (frequency x injected current) and number of spikes x injected current (bottom) are plotted for each age group of males (C., blue, left) and females (D., green, right). Box and whisker plots show slopes of F/I curve (E) and number of spikes at 400 pA current injection across ages in males (blue) and females (green). Red dashed lines show means. Bonferroni corrected t-test: males and females differ, ***p<0.001; *a* – 18 and 24 month males differ from 12 month males, p<0.05; *b* – 18 month males differ from 4 month males, p<0.05; # - 24 month males differ from 4 month males, p=0.057.

**Figure 4.**
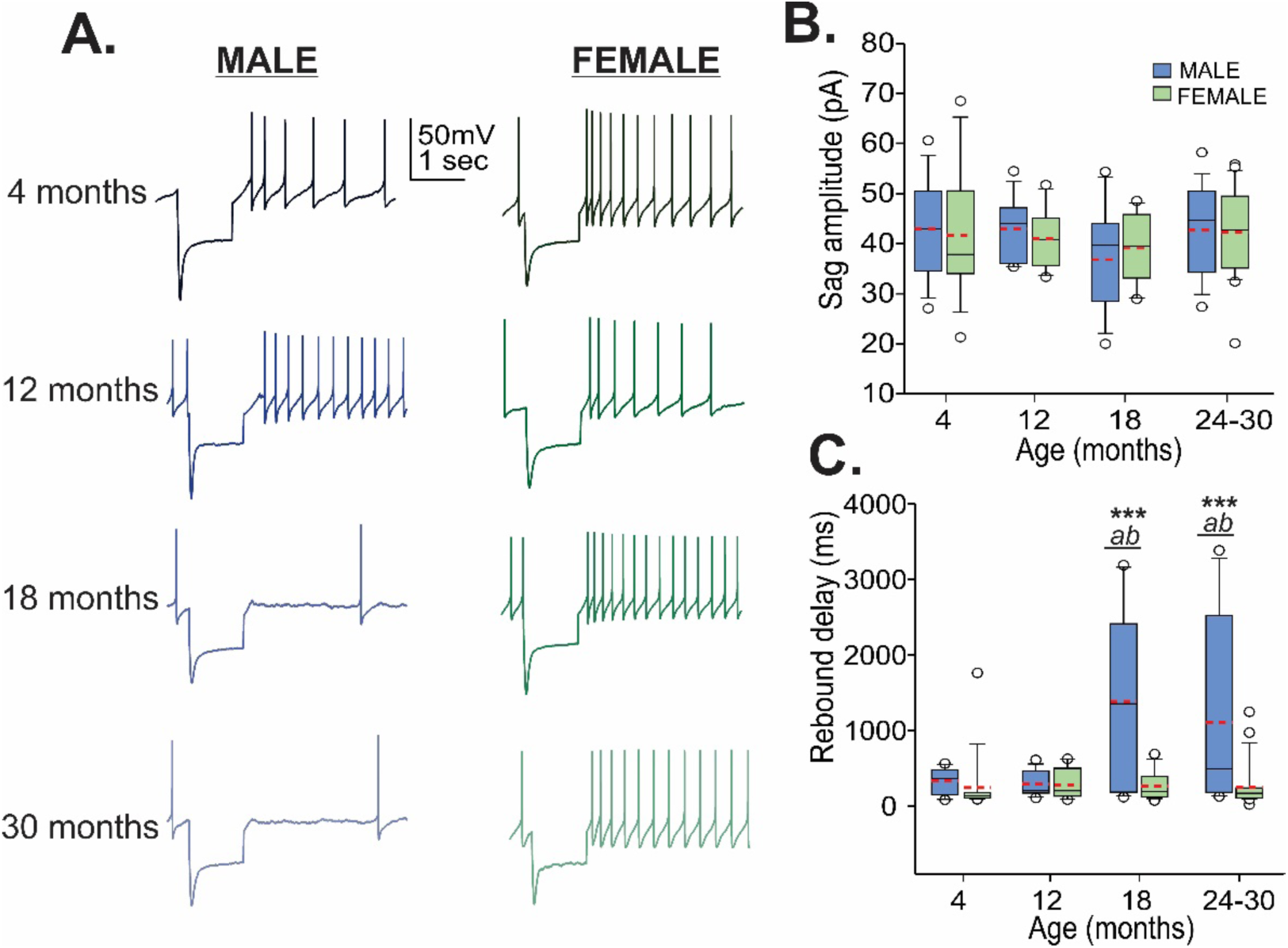
Sag and post inhibitory rebound spiking across ages in males and females. A. Representative traces of hyperpolarizing current injections at 4, 12, 18, and 24-30 months from males (left, blue) and females (right, green). Box and whisker plots show sag amplitude (B) across ages (males, blue; females, green) and time delay until rebound spike (C). Red dashed lines show means. Bonferroni corrected t-test: males and females differ, ***p<0.001; *a* – 18 and 24 month males differ from 12 month males, p<0.01; *b* – 18 and 24 month males differ from 4 month males, p<0.02

**Figure 5.**
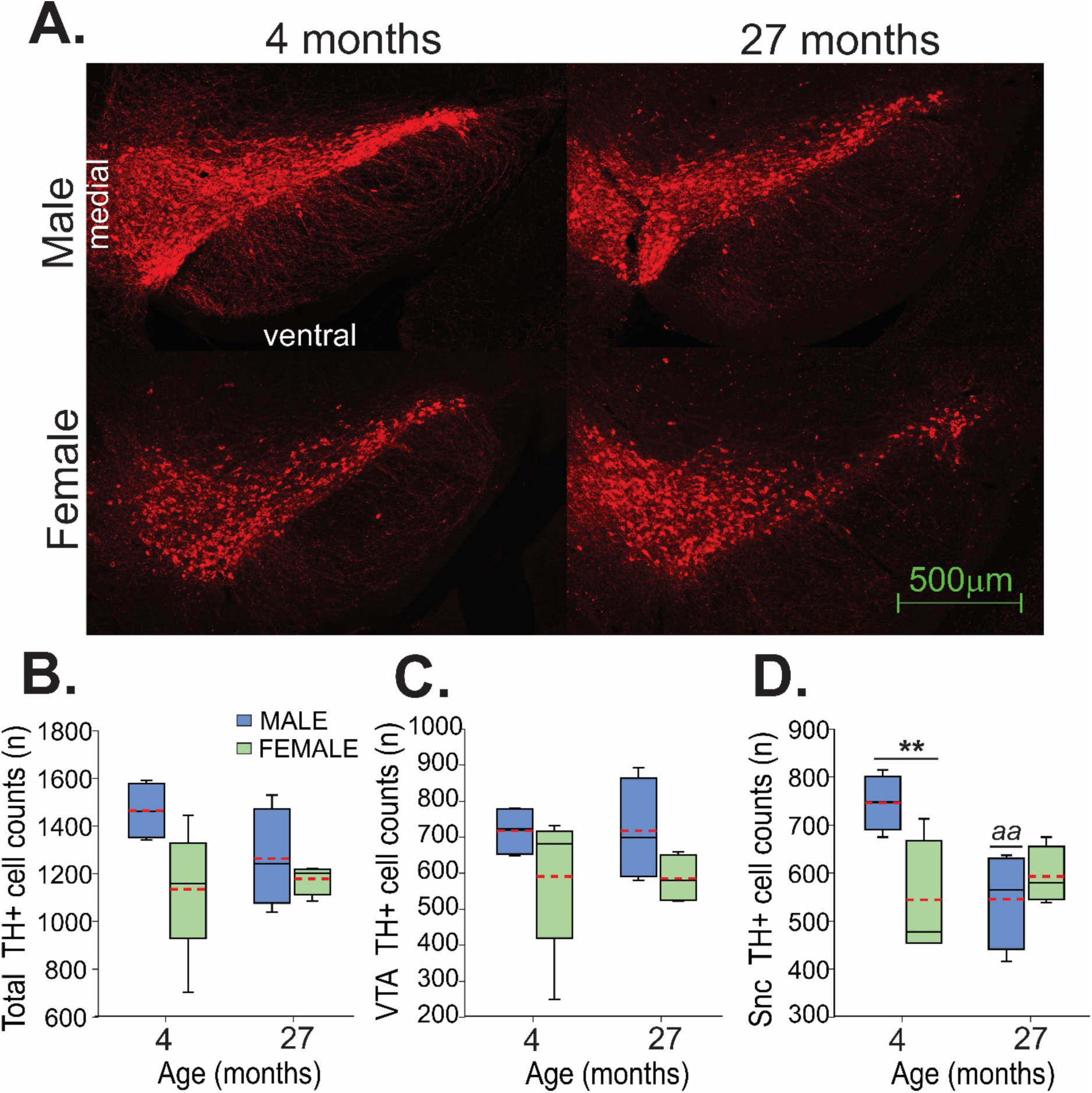
Semi-quantitative analysis of tyrosine hydroxylase positive neurons. A. Representative images at -3.08mm from bregma from a 4-month-old male and female (left) and 27 month old male and female (right). B. Box and whisker plot showing total TH positive cell counts across the three levels quantified (−3.08, -3.28, -3.52 from bregma) in 4 month and 27-month-old males (blue) and females (green). Red dashed lines show means. Bonferroni corrected t-test: males and females differ, *p<0.05; old males differ from young males, a – p<0.01.

**Figure 6.**
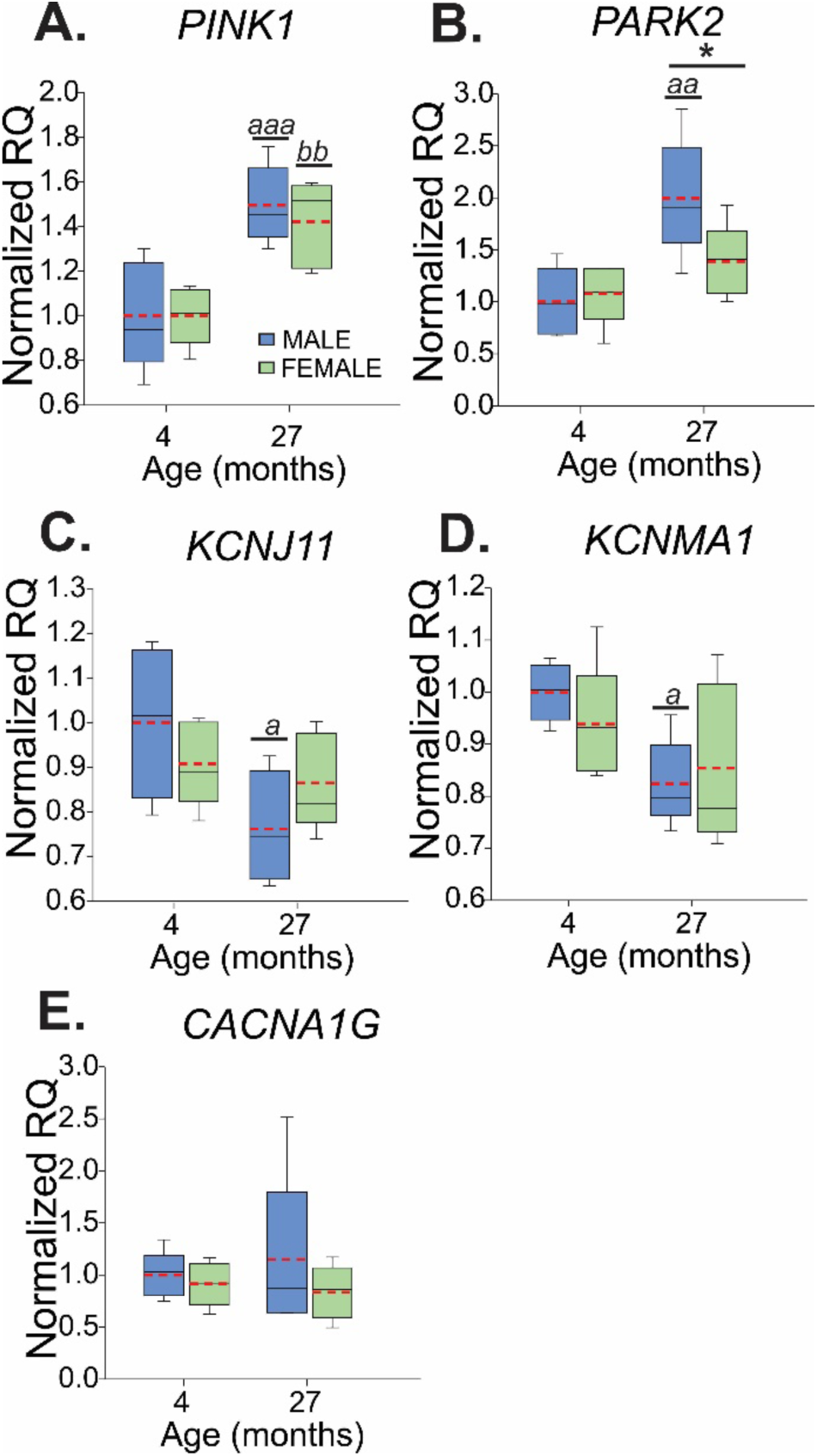
Gene expression changes between males (blue) and females (green) with age for PTEN induced putative kinase (*PINK1*, A), parkin (*PARK2*, B), K-ATP channels (*KCNJ11*, C), BK channels (*KCNMA1*, D), and T-type calcium channels (*CACNA1G*, E). Box and whisker plots show relative quantification (RQ) for male (blue) and female (green) animals at 4 months and 27 months of age. Rd dashed lines indicate means. Bonferroni t-test: Males differ from females *p<0.05; old males differ from young males, a - p<0.05, aa - p<0.005, aaa - p<0.001; old females differ from young females, bb - p<0.005

Males showed a robust age-current interaction for number of spikes (F_57,1672_=21.142, p<0.001). 18-month old males showed decreases in spikes evoked by a 200 pA current injection and 24-30 months old males showed decreases by a 300 pA injection. Females did not show any change in the number of spikes across ages. Overall, females showed higher numbers of spikes at all ages compared with males (F_1,166_=7.378, p=0.007) and a higher maximum number of spikes (F_1,166_=6.666, p=0.011). Similarly, males 18 months and older showed fewer spikes for the highest current injection (400pA) compared with females (F_3,166_=3.717, p=0.013, Fig 3F). No bimodal distribution was observed for the number of spikes from dopamine neurons recorded from males; almost 90% of cells from males 18 months and older showed spike numbers below the average for young males at a 400pA current injection. This decrease in spikes at higher current injections in older males seems to indicate a greater tendency of dopamine cells to enter depolarization block than in younger males or females at any age.

### Dopamine neurons from males 18 months and older, but not females, show a longer rebound spike delay

Dopamine neurons are characterized by I_H_, which can be measured in current clamp as a large sag during hyperpolarizing current injections (Dufour et al., 2014). We next assessed I_H_ and rebound firing properties in dopamine cells by delivering a hyperpolarizing current injection for 1 second, bringing the potential at the end of the sag to approximately –80 mV (−83.73±1.007 mV across all cells). Sag and rebound spiking were recorded from animals at 4 months (n=13 from 4 males; n=15 from 3 females), 12 months (n=13 from 5 males; n=12 from 5 females), 18 months (n=17 from 4 males; n=17 from 3 females) and 24-30 months (n=16 from 8 males; n=21 from 7 females). No change was detected in sag amplitude across ages and between sexes, suggesting no change in I_H_. This agrees with our previous findings of no change in I_H_ with age in males, measured in voltage clamp (Branch et al., 2014). However, there was a robust age-sex interaction detected for rebound spike delay (F_3,116_=5.177, p=0.002). At ages 18 months and older, males show a 483% greater average than females of the same age for rebound spike delay. As with the CV_ISI_, a similar proportion of dopamine cells from males ages 18 months and older showed rebound spike delays in the range of the young males, 35.3% of the cells from the 18-month-old group and 50% of the cells from the 24-30 month age group (Fig S3). Further analysis showed that long rebound spike delays in cells old males correlated with increasing CV_ISI_ and lower firing frequencies (Fig S4). At 18 months, those “healthy” dopamine cells were recorded from each of the animals in that group, while at 24-30 months those cells came from 62.5% of the animals.

### Males show loss of midbrain dopamine neurons with age, while females do not

Based on the electrophysiological findings, at least 60% of dopamine neurons recorded from males aged 18 months and older showed robust deficits in pacemaking and rebound spiking, while females showed very few deficits. We next examined how the population of dopamine neurons in the region was affected by aging. A semi-quantitative analysis of TH positive neurons in the midbrain was performed in mice at 4 months of age (n=4, males; n=5 females) and 27 months of age (n=4, males; n=4 females). TH positive cell counts were compiled from coronal slices containing the ventral region of the midbrain (SN and VTA) at -3.08 mm, -3.28 mm, and - 3.52 mm from bregma. In the whole ventral midbrain, females had apparently fewer dopamine neurons than males (F_1,13_=4.997, p=0.044), approximately 84.8% of the cells counted in males overall (combining both age groups). Young females had 77.5% of the cells of young males, as previously observed (Dewing et al., 2006). In our sampling, no differences were detected between age and sex in the VTA, however fewer dopamine neurons were observed in the SN only in males with age (interaction: F_1,13_=8.046, p=0.014). Old males showed a 26.9% loss of dopamine neurons in the SN compared with young males.

### PINK1 and PARK2 expression increases with age in the SN, primarily in males

To test if alterations in mitochondrial function and calcium signaling play a role in the deficits in firing activity observed in males with age and the differences with females, qPCR was performed to evaluate the expression of mRNA for ion channel and Parkinson’s-related genes: *PARK2, PINK1, KCNJ11, CACNA1G* and *KCNMA1*. SN tissue was harvested from 4-month-old (n=5 males; n=5 females) and 27-month-old mice (n=5 males; n=4 females). In old mice, *PINK1* expression increased by 146% (age: F_1,16_=29.636, p<0.001) and *PARK2* expression increased by 163% (age: F_1,16_=13.268, p=0.002). Additionally, for *PARK2* there was a trend toward an age-sex interaction (F_1,16_=3.667, p=0.074), with males having an increase in expression of 185% (Bonferroni t-test, p=0.001) and females having an increase of 139% (Bonferroni t-test, p=0.24). Decreases were observed in expression of mRNA for the K-ATP channel subunit (*KCNJ11*) between young and old animals (F_1,15_=5.717, p=0.03). Overall there was a 14.8% decrease in K-ATP mRNA expression between young and old mice, with males showing a 23.8% decrease (Bonferroni t-test, p=0.014) and females showing no change (Bonferroni t-test, p=0.6). A decrease in mRNA for BK channels (*KCNMA1*, “big” conductance calcium activated potassium channels) was observed with age (F_1,16_=7.286, p=0.016). Comparing young and old animals alone, there was a 13.4% decrease with age; 17.6% in males (Bonferroni t-test, p=0.02) and 8.9% in females (Bonferroni t-test, p=0.23). No significant differences were detected in expression of T-type calcium channels (*CACNA1G*).

## DISCUSSION

Here we report the first in-depth characterization of dopamine neuron firing parameters across the lifespan in both male and female mice. We observed that dopamine neurons from male mice begin showing signs of deterioration in firing activity by 18 months of age, while dopamine neurons from females were more resilient throughout the lifespan. Impairments observed in dopamine neurons from males included decreases in firing rate, disrupted pacemaking, increased propensity to enter depolarization block, and prolonged delays in rebound spiking. Impairments in rebound spiking were correlated with decreased firing rate and pacemaking deficits, suggesting disruption in a similar underlying process (Fig S4). Interestingly, the depolarization block seemed to be inherently present in the majority of cells from males 18 months and older. In addition, a semi-quantitative analysis suggested a slight decrease in dopamine neuron number in the SN with age only in males. Accompanying those changes were increases in *PINK1* and *PARK2* mRNA expression with age, suggesting an increased response to mitochondrial stress. Males and females only showed slight decreases in locomotor activity across ages with no obvious sex effect (Fig S1). Overall, we interpret the age-dependent alterations in dopamine neuronal properties in the males as possibly contributing a vulnerability to further insults rather than a disease process *per se*, while dopamine neurons from females show resilience to these effects of normal aging.

### Sex divergence in dopamine neuron action potential generation with age

Firing properties of SN dopamine neurons have only been lightly studied in females and not at all studied in aged female mice. We identified robust phenotypical differences in SN dopamine neuron firing properties between aged male and female mice. At the youngest age tested (4 months), SN dopamine neurons from females exhibited a larger spike height and faster action potential kinetics compared to males. In fact, the only change in neuronal firing properties that occurred in females was a linear decrease in spike height and kinetics with age. By 24 months it appeared that individual males segregated into two populations that could be designated as “aged-healthy” and “aged-impaired”. Indeed, this phenomenon has been widely documented with other phenotypes in rodents over 24 months of age (Menard et al., 2015; Soontornniyomkij et al., 2010). We also observed that young females had fewer dopamine neurons in the midbrain compared with young males, comparable to other reports (Dewing et al., 2006). In addition, young female mice also showed higher basal locomotor activity than males, which has also been previously observed (Van Swearingen, Walker, & Kuhn, 2013). Both sexes showed a similar modest decline in spontaneous locomotion with age; these experiments were conducted in the dark which is the active phase in rodents (Fig S1). Consistent with this, Fischer et al. (2016) previously reported similar decreases in animals tested in the dark phase, however in the light phase they only observed a decrease in locomotor activity in males at 28 months of age. Therefore, while declining firing properties of dopamine neurons from males were not accompanied by mass cell death or a sexually divergent effect in spontaneous locomotion, they could instead reflect an intrinsic vulnerability to additional insults.

In the context of Parkinson’s disease, evidence from lesion-based models of Parkinson’s suggests biological differences in SN dopamine neurons from females compared with males. Administration of MPTP (1-methyl-4-phenyl-1,2,3,6-tetrahydropyridine) or 6-OHDA (6-hydroxydopamine), toxins which target mitochondrial respiration (Glinka, Gassen, & Youdim, 1997; Przedborski, Tieu, Perier, & Vila, 2004), produce a greater loss of SN dopamine neurons in males compared with females (for review see Gillies et al., 2014). This is also true in aged rats, where following toxin exposure females experience far less of a loss of dopamine neurons compared with males (Tamas, Lubics, Szalontay, Lengvari, & Reglodi, 2005). Genes escaping X-linked inactivation have failed to account for the decreased susceptibility of females to Parkinson’s (Sharma et al., 2017), but ovarian steroid studies have accounted for the resilience of females to oxidative damage. In whole brain samples, young females showed lower levels of oxidative stress than males, which was absent in animals that were either ovariectomized or aged (Gaignard et al., 2015). Indeed, human clinical studies have shown an increased risk of developing Parkinson’s after hysterectomy with removal of both ovaries (Benedetti et al., 2001). While mice do not formally enter menopause like humans, by 18 months of age they are reproductively senescent and have very low plasma levels of estrogen and progesterone (Lu, Hopper, Vargo, & Yen, 1979). As the degeneration of dopamine neurons is a slow process, it is possible the months of cycling hormones can afford females with the prolonged protection needed to delay the decline of dopamine neuron function.

### Vulnerability of SN dopamine neurons and risk of Parkinson’s disease

Previous work from our lab using a progressive mouse model of Parkinson’s disease through impairment of mitochondrial function found that deficits in multiple electrical parameters, including pacemaking, preceded degeneration of dopamine cells (Branch et al., 2016). Aging is thought to create a pre-Parkinsonian environment in dopamine cells through oxidative damage, mitochondrial dysfunction, and inflammation, thereby promoting neurodegeneration following exposure to additional insults such as toxins or infections (Collier, Kanaan, & Kordower, 2017). Consistent with our findings, previous studies have described moderate decreases in SN dopamine cell number in males of several species including humans (McGeer, Itagaki, Akiyama, & McGeer, 1988), non-human primates (Collier et al., 2017), rats (Gao et al., 2011), and mice (Wey et al., 2012). In Parkinson’s, motor deficits are largely undetectable until at least 50% of dopamine neurons in the SN are lost (Cheng, Ulane, & Burke, 2010). Therefore, the modest reduction in spontaneous locomotor activity we observed is not unexpected as the age-related cell loss we observed was well short of 50%. Dopamine neurons of the SN are noteworthy for their vulnerability to oxidative stress (Duda et al., 2016) as opposed to dopamine neurons of the VTA, which are more resistant to neurodegeneration (Damier, Hirsch, Agid, & Graybiel, 1999). As autonomous pacemakers, SN dopamine cells depend on large, well-controlled calcium oscillations for their continuous firing activity (Puopolo, Raviola, & Bean, 2007), which creates great energy demands for the cell (Duda et al., 2016). Mitochondrial damage, produced for example through age-related accumulation in oxidative stress, can increase intracellular calcium concentrations and produce adverse consequences such as excitotoxicity, apoptosis, and activation of detrimental enzymatic cascades (Mattson, 2007).

The increase we observed in *PINK1* and *PARK2* expression, particularly in males, could suggest age-related damage to mitochondria and possibly increases in intracellular calcium. *PINK1* encodes PTEN induced putative kinase 1, which accumulates on damaged mitochondria and works with parkin (encoded by *PARK2*), the E3 ubiquitin ligase, in facilitating mitochondrial degradation (Srivastava, 2017). Notably, mutations in *PINK1* and *PARK2* have been identified in autosomal recessive forms of early onset Parkinson’s disease, causing accumulation of damaged mitochondria and leading to dopamine neuron degeneration. Moreover, high intracellular calcium levels can induce an increase in *PINK1* expression (Gomez-Sanchez et al., 2014). Interestingly, *PINK1* mRNA is upregulated in SN dopamine neurons in brains of old male human subjects, but not in females (Cantuti-Castelvetri et al., 2007).

### Participation of ion channels in the effects of age in males

Disruption in dopamine neuron function leading to increased intracellular calcium has been long hypothesized to underlie decrements associated with aging and neurodegenerative disorders (Alzheimer’s Association Calcium Hypothesis, 2017). A continuing rise in calcium with increasing age would not only affect intracellular signaling cascades but would also alter ion channel function. From our cross-sectional sampling, action potential waveforms from male mice showed decreases in spike height and prolonged kinetics at the oldest ages tested. However, our half-width data showed a biphasic effect across the lifespan in males, with decreases in half-width by 12 months followed by a progressive decline with age. Our qPCR data indicated a decrease in transcript for *Kcnma1*, the gene that codes for the “big conductance” calcium activated potassium (BK) channel subunit in aged males. Studies in CA1 pyramidal cells found increases in activity of calcium activated potassium channels, due to increased intracellular calcium with age, resulting in increases in kinetics and afterhyperpolarization (Gant & Thibault, 2009). However, SN dopamine neurons are far more sensitive to calcium than other neuron types due to their low calcium buffering capacity (Duda et al., 2016). Given that signs of oxidative stress are already present in dopamine neurons from animals no older than 1 month (Guzman et al., 2018), it is possible that increases in intracellular calcium may be present at 12 months and affect action potential parameters. Decreased BK channel activity with increasing age would be expected to widen the half-width (Kimm, Khaliq, & Bean, 2015), which is consistent with our observations in males aged 18 months and older.

We also observed decreased mRNA expression of *Kcnj11*, the gene that codes for Kir6.2/K-ATP channels. These channels act as energy sensors in dopamine neurons and have been proposed to play a role in the development of neurodegeneration, however their activation has paradoxically been shown to increase bursting (Schiemann et al., 2012). Dopamine cells from aged males also exhibited delayed rebound spiking, which could indicate decreased function of T-type calcium channels (Evans, Zhu, & Khaliq, 2017). However, we did not detect a change in mRNA transcript levels for *Cacna1g*, the gene coding for Cav3.1 subunits. We previously showed decreased nimodipine-sensitive (presumably L-type calcium) currents in aged male mice, but also similarly no difference in mRNA transcript levels (Branch et al., 2014). It is therefore possible that these channels are regulated by membrane trafficking or at the level of translation. Alternatively, their decreased function could indicate greater inactivation produced in response to elevated levels of intracellular calcium (Simms & Zamponi, 2014), which could also explain the decrease in afterhyperpolarization potential we observed in neurons from aged males.

## Conclusion

The results presented here demonstrate the existence of functional electrophysiological differences between male and female dopamine neurons of the mouse SN in response to aging. Dopamine neurons from female mice were resistant to detrimental consequences of aging, which also included less dopamine cell loss in the SN. The results provide converging evidence that SN dopamine neurons in males are more vulnerable to the effects of aging, paralleling epidemiological evidence indicating that males have nearly twice the incidence of Parkinson’s as females.

## EXPERIMENTAL PROCEDURES

### Animals

Male and female C57BL/6 mice were obtained from the National Institute on Aging aged rodent colony and housed in a humidity and temperature-controlled facility under a reverse 12:12 light-dark cycle (lights off 0900) with food and water available *ad libitum*. Animals were examined at 4 different ages: 4 months, 12 months, 18 months and 24-30 months. All experiments were reviewed and approved by the Institutional Animal Care and Use Committee at the Oklahoma Medical Research Foundation.

### Ex vivo electrophysiology

Mice were deeply anesthetized with isoflurane before rapid dissection of the brain, which was immediately placed in an ice-cold oxygenated cutting solution containing the following in mM: 250 sucrose, 26 NaHCO_3_, 2 KCl, 1.2 NaH_2_PO_4_, 11 glucose, 7 MgCl_2_, 0.5 CaCl_2_. Coronal slices (200 µm) containing ventral midbrain were collected using a VF-200 Compresstome (Precisionary, Cambridge, MA, USA). Slices were incubated at 34°C for 20 minutes, after which they were incubated at room temperature in an artificial cerebrospinal fluid (aCSF) solution containing the following in mM: 126 NaCl, 2.5 KCl, 1.2 NaH_2_PO_4_, 26 NaHCO_3_, 11 glucose, 1.3 MgCl_2_, 2 CaCl_2_ and with 0.01 MK-801. During recording, slices were superfused with aCSF at 34-36°C with a flow rate of 2 ml/min in the presence of 100 µM picrotoxin and 10 µM DNQX to antagonize any spontaneous GABAergic and glutamatergic synaptic activity that could influence the cell firing parameters that were measured.

Dopamine neurons were visualized using an upright microscope (Nikon, Melville, NY, USA) a patch clamped using thin wall glass pipettes (1.5-3 MΩ, World Precision Instruments, Sarasota, FL, USA) filled with internal solution containing the following (in mM): 142 K-gluconate, 2 KCl, 4 MgCl_2_, 10 HEPES, 0.5 EGTA, 4 ATP-Mg, 0.5 Na-GTP, 280-300 mOsm, pH adjusted to 7.2 to 7.4 with KOH. In addition, 0.05% neurobiotin (Dufour et al., 2014) was added to the patch pipette to allow for *post hoc* verification of the dopamine identity of neurons recorded, through immunolabeling for tyrosine hydroxylase (see below). Current clamp recordings were made with a Multiclamp 700B amplifier (Molecular Devices, Sunnyvale, CA, USA), filtered at 4 kHz and digitized at 50 kHz. Data were collected using Axograph software (www.axograph.com). Voltages were corrected for a liquid junction potential of 11.1 mV between the internal and external solution. Electrophysiological data were analyzed with IgorPro (Wavemetrics, Lake Oswego, OR, USA).

### Immunohistochemistry

Animals (n = 4-5, per age and sex) were deeply anesthetized with tribromoethanol (250 mg/kg, i.p.) and perfused by intracardiac puncture with 20 ml of phosphate-buffered saline (PBS; 0.9% NaCl in 50 mM phosphate buffer, pH 7.4), followed by 20 ml of 4 % paraformaldehyde (PFA) solution in 0.1 M phosphate buffer (PB, pH 7.4). The brain was removed, post-fixed overnight in 4 % PFA solution at 4 °C, washed in PBS and immersed in 30% sucrose at 4°C for 3 days. After this, the brain was embedded in OCT, stored at -80 °C and processed as described below.

#### Cryostat cutting

Consecutive coronal sections (40 µm thick) were collected throughout the whole extension of the ventral midbrain containing the VTA and the SN, between stereotaxic coordinates 2.9 to 3.88 posterior to Bregma (Franklin & Paxinos, 2013). The sections were collected in wells containing PBS for further processing.

#### Tyrosine hydroxylase (TH) immunolabeling

Polyclonal rabbit anti-tyrosine hydroxylase antibody (Millipore-Sigma, Burlington) was used as our primary antibody and goat anti-rabbit Alexa Fluor 594 (Abcam, Cambridge) was used as secondary antibody. Normal goat serum (Jackson ImmunoResearch, West Grove) was used in all blocking steps.

Brain sections were equilibrated to room temperature (RT) for 30 minutes, rinsed in PBS and incubated in a blocking solution (BS; 5% normal goat serum in 0.2% PBST, phosphase buffer saline tris) for 4 hours at RT in a platform shaker at low speed. Then, sections were incubated overnight with primary TH antibody (1:750 dilution in 0.5% normal goat serum in PBS) at 4 °C with gentle shaker agitation. After rinsing with 0.2% PBST, sections were incubated with the secondary antibody Alexa Fluor 594 (1:200 dilution in PBS) for 2 hours at RT and protected from light with foil. After rinsing with PBST and PBS, sections were mounted onto glass slides using Shur/Mount Aqueous (EMS, Hatfield). Coverslips were applied and sealed with nail polish.

For neurobiotin-filled cells, brain slices from electrophysiology experiments were fixed in 4% PFA overnight and then processed for TH immunolabeling as described above. Natural streptavidin protein Dylight 488 (1:500 dilution in PBS, Abcam, Cambridge) was added to label neurobiotin. 100 ng/ml DAPI in 0.2% PBST (1.5 hrs. incubation) was added in the last rinse after secondary antibody incubation. This was followed by three rinse steps (1-hour, 30 min, 30 min) with PBST to remove excess staining. Brain slices were then mounted on glass slides as described above.

#### Image acquisition and cell counting

Images were obtained using a Zeiss LSM-710 confocal microscope with a 10X magnification objective. High resolution images were stitched to include the range of ventral midbrain containing the VTA and SN. Semi-quantitative cell counting was done using Image J software.

### qPCR

The SN was dissected from mice 4 months old (5 males, 5 females) and 30 months old (5 males and 5 females). Samples were immediately snap-frozen in liquid nitrogen and stored at -80°C. Total RNA was extracted from frozen tissue using RNeasy plus mini kit (Qiagen, CA, USA). The amount of total RNA was quantified using Nanodrop 2000 spectrophotometer and RNA quality and purity was evaluated using Agilent 2200 TapeStation system (Agilent Technologies, CA, USA). One sample failed quality control leaving 4 mice in the aged female group. Gene expression levels were assayed by qPCR, as previously described (Mangold et al., 2017; Masser et al., 2014). Briefly, cDNA was synthesized from 80 ng RNA using Quantabio qScript cDNA Sythesis Kit. Pre-designed probe and primer fluorogenic exonuclease assays (TaqMan, Life Technologies, Watham, MA, Table S1) were used to perform qPCR on the QuantStudio™ 12K Flex Real-Time PCR System (Applied Biosystems). Relative gene expression (RQ) was normalized to endogenous control GAPDH using the 2^−ΔΔCt^ method in the Expression Suite v1.0.3 software.

### Statistical analysis

Data are represented in box and whisker plots, to show distributions of data, or line graphs with means ± standard error of the mean. Data were analyzed by two-way ANOVA (factors: sex, age) and two-way repeated measures (RM) ANOVA, followed by Bonferroni t-test post hoc (Sigma Stat, San Jose, CA, USA). Where indicated, Bonferroni corrected t-tests were performed. Alpha was set to 0.05.

## ACKNOWLEDGEMENTS

We would like to thank Julie Crane, Justin Willige and Ben Fowler from the imaging core facility at the Oklahoma Medical Research Foundation for their assistance with the immunostaining and confocal imaging. We would also like to thank Dr. Ishita Parikh for isolating the RNA and preparing cDNA from the tissue samples. This work was supported by National Institute on Aging/NIH grants R01 AG052606 and P30 AG050911, as well as funds from the Presbyterian Health Foundation and the Oklahoma Center for Adult Stem Cell Research (OCASCR). The authors declare no conflicts of interest.

## AUTHOR CONTRIBUTIONS

RDH performed all electrophysiology, collected tissue for qPCR, quantified images from immunocytochemistry, designed the experiments and wrote the paper. SDL collected tissue for qPCR, processed images from immunocytochemistry, performed locomotion studies and helped with manuscript preparation. SO and WMF performed qPCR experiments. MJB helped with experimental design and writing the paper.

## SUPPORTING INFORMATION

**Table S1.** Taqman gene assays used for qPCR analysis.

| **Gene/Gene ID** | **Gene Name** | **Taqman Gene Expression assay ID** |
| --- | --- | --- |
| *Park2* | Parkinson disease (autosomal recessive, juvenile) 2, parkin | Mm01323528_m1 |
| *Pink1* | PTEN induced putative kinase 1 | Mm00550827_m1 |
| *Kcnj11* | potassium inwardly rectifying channel, subfamily J, member 11 | Mm00440050_s1 |
| *Cacna1g* | calcium channel, voltage-dependent, T type, alpha 1G subunit | Mm00486572_m1 |
| *Kcnma1* | potassium large conductance calcium-activated channel, subfamily M, alpha member 1 | Mm01268569_m1 |
| *Gapdh* | Glyceraldehyde-3-Phosphate Dehydrogenase | 4352661 |

**Figure S1:**
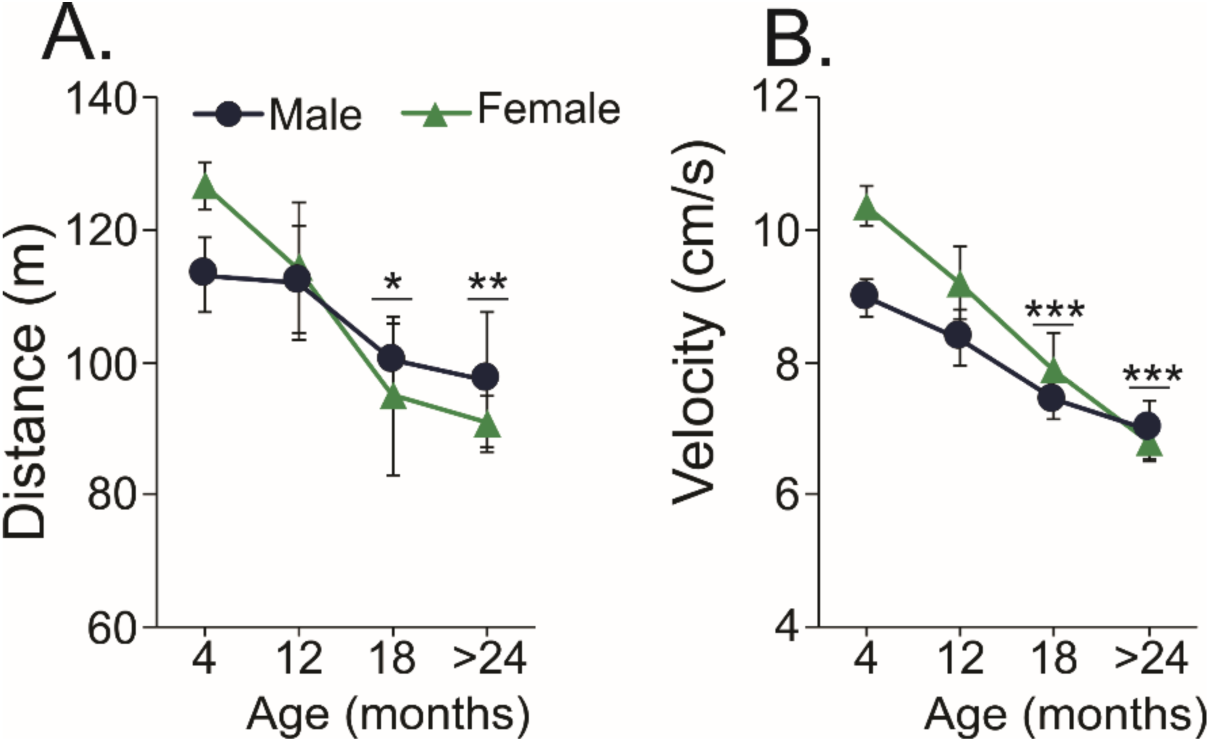
Assessment of open field locomotion across age. Spontaneous locomotor activity was assessed for 30 minutes in animals across ages 4 months (n=15 males; n=13 females), 12 months (n=11 males; n=8 females), 18 months (n=10 males; n=8 females), and 24-30 months (n=7 males; n=8 females). Mice were placed in open field locomotor chambers (44.5 × 44.5 cm; Columbus Instruments, Columbus, OH, USA) equipped with infrared photosensors. Quantification was performed using Autotrack 5.5 software (Columbus Instruments). Distance traveled and average velocity during movement both decreased by 18 months of age in males and females (distance: F_3,72_=5.771, p=0.001; velocity: F_3,72_=20.41, p<0.001). Females overall showed higher movement velocities than males (F_1,72_=4.835, p=0.031), which was most noticeable in the youngest age group. A. Total distance traveled by males (blue circles) and females (green triangles). B. Average velocity of mice during movement. Bonferroni corrected t-test: 18 or 24 months differs from 4 months, *p<0.05, **p<0.01, ***p<0.001.

**Figure S2:**
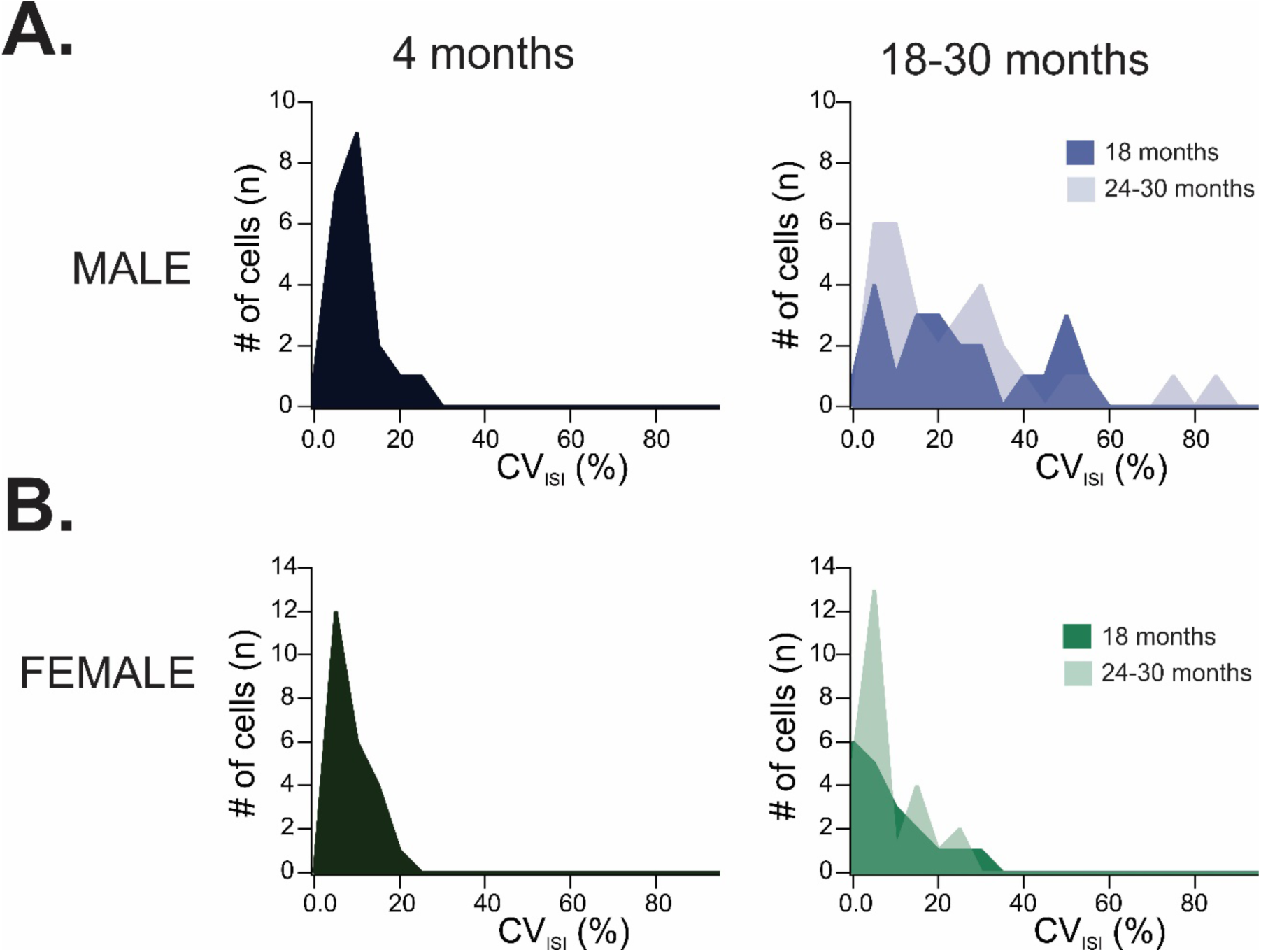
Histograms depicting distribution of CV_ISI_ at 4 months and 18-30 months for males (A, blue, top) and females (B, green, bottom).

**Figure S3:**
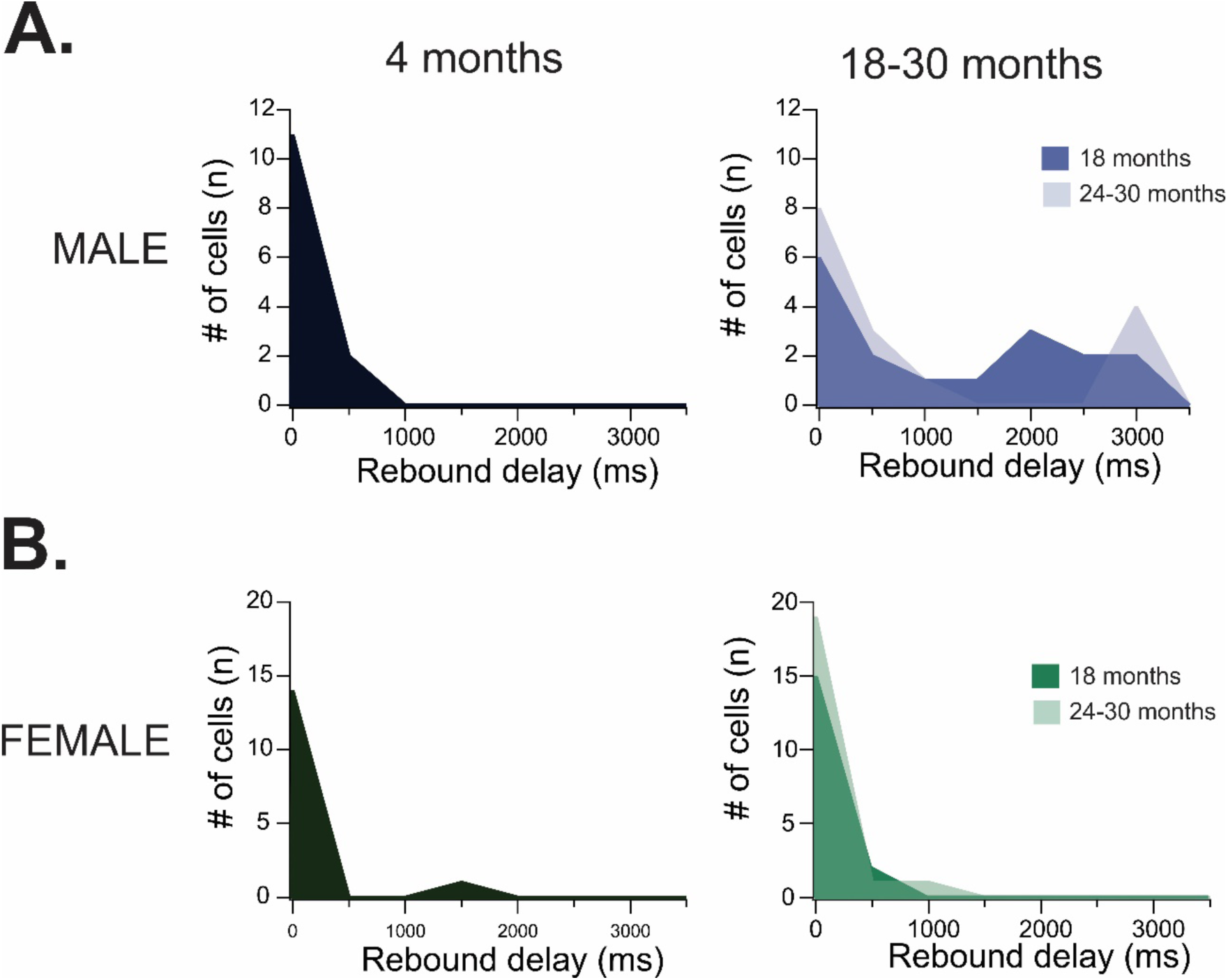
Histograms depicting distribution of rebound spike delay at 4 months and 18-30 months for males (A, blue, top) and females (B, green, bottom).

**Figure S4:**
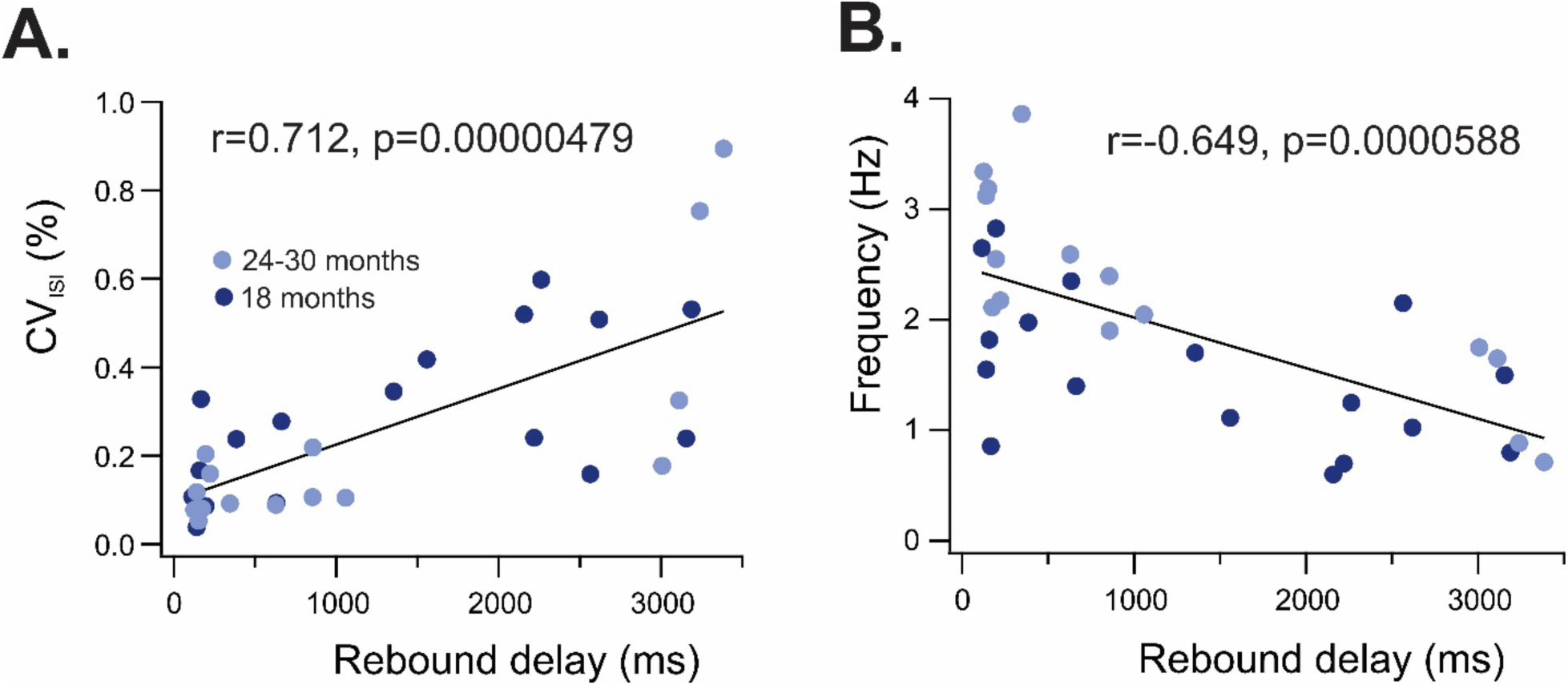
Pearson correlations of rebound spike delay and either CV_ISI_ (A) or firing frequency (B) in males ages 18-30 months. Cells from 18-month-old males are indicated by dark blue circles and cells from 24-30-month-old males are indicated by light blue circles.

